# On standardization of controls in lifespan studies

**DOI:** 10.1101/2023.08.17.552381

**Authors:** Olga Spiridonova, Dmitrii Kriukov, Nikolai Nemirovich-Danchenko, Leonid Peshkin

**Affiliations:** Department of Systems Biology, Harvard Medical School, Boston, MA 02115, USA

**Author notes:** These authors contributed equally (Olga Spiridonova, Dmitrii Kriukov).

## Abstract

The search for interventions to slow down and even reverse aging is a burgeoning field. The literature cites hundreds of supposedly beneficial pharmacological and genetic interventions in model organisms: mice, rats, flies and worms, where research into physiology is routinely accompanied by lifespan data. Naturally the negative results are more frequent, yet scientifically quite valuable if analyzed systematically. Yet, there is a strong “discovery bias”, i.e. results of interventions which turn out not to be beneficial remain unpublished. Theoretically, all lifespan data is ripe for re-analysis: we could contrast the molecular targets and pathways across studies and help focus the further search for interventions. Alas, the results of most longevity studies are difficult to compare. This is in part because there are no clear, universally accepted standards for conducting such experiments or even for reporting such data. The situation is worsened by the fact that the authors often do not describe experimental conditions completely. As a result, works on longevity make up a set of precedents, each of which might be interesting in its own right, yet incoherent and incomparable. Here we point out specific issues and propose solutions for quality control by checking both inter- and intra-study consistency of lifespan data.

## Introduction

In the fascinating realm of aging biology, a critical challenge emerges as we delve into the heart of various studies. The enigma lies in the stark inconsistency that surfaces when we meticulously gather control data across these studies – data from those untouched by interventions, a true baseline to measure against. This inter-study inconsistency becomes evident if we compile the control data (i.e. animals that have not been subject to interventions) across studies. We focus on male [C57BL/6J](https://www.jax.org/strain/000664) mice - the most widely used inbred strain in biology of aging. According to [JAX® Mice & Services](https://www.jax.org/strain/000664) / [Mouse Phenome Database](https://phenome.jax.org/strains/7)^9^, the median lifespan of C57BL/6J males is 894-901 days, another source^1^ indicates a life expectancy of 878 ± 10 days. Our own literature review (Fig. S1) suggests that in various studies the median control lifespan of C57BL/6J animals varies from 600 to 980 days. It is impossible to reliably determine the causes of lifespan variability as it is affected by diet, cage density, temperature, light cycle, etc. In addition, the C57BL/6J line has a long breeding history, and mice taken from different institutions have genetic and physiological features that affect the results of different tests^2^. The road to standardization and quality control of mouse control survival data must necessarily be traversed by someone. In this short article, we offer to go through it with us.

## Results

To further study the life expectancy standard for male C57BL/6J we selected ‘meta-controls’ - control data from several papers where C57BL/6J mice came from two sources: the National Institute of Aging^3–5^ and Jackson Laboratories^6,7^ and the origins and conditions of the control animals were described in sufficient detail. Across these studies, the median longevity varies between 800 and 970 days -- less than in studies with C57BL/6J males in general. Crucially, even this range far exceeds the typical difference between the median lifespan of a “successful” intervention and control groups, which generally does not exceed 15%. Indeed, when we looked at some of the best known studies on experimental life extension in C57BL/6J males - we found that in most cases the median lifespan of the experiment, although significantly longer than that of respective control - still did not exceed the longest median lifespan across the ‘meta-controls’ ^6^ (Fig. 1) - one notable exception being the rapamycin^8^. Each dataset also was tested with our newly developed extra-mortality test (see Supplement) showing if the given dataset contains implausible instantaneous increase in mortality. A sudden burst of deaths suggests latent issues.

**Figure 1.**
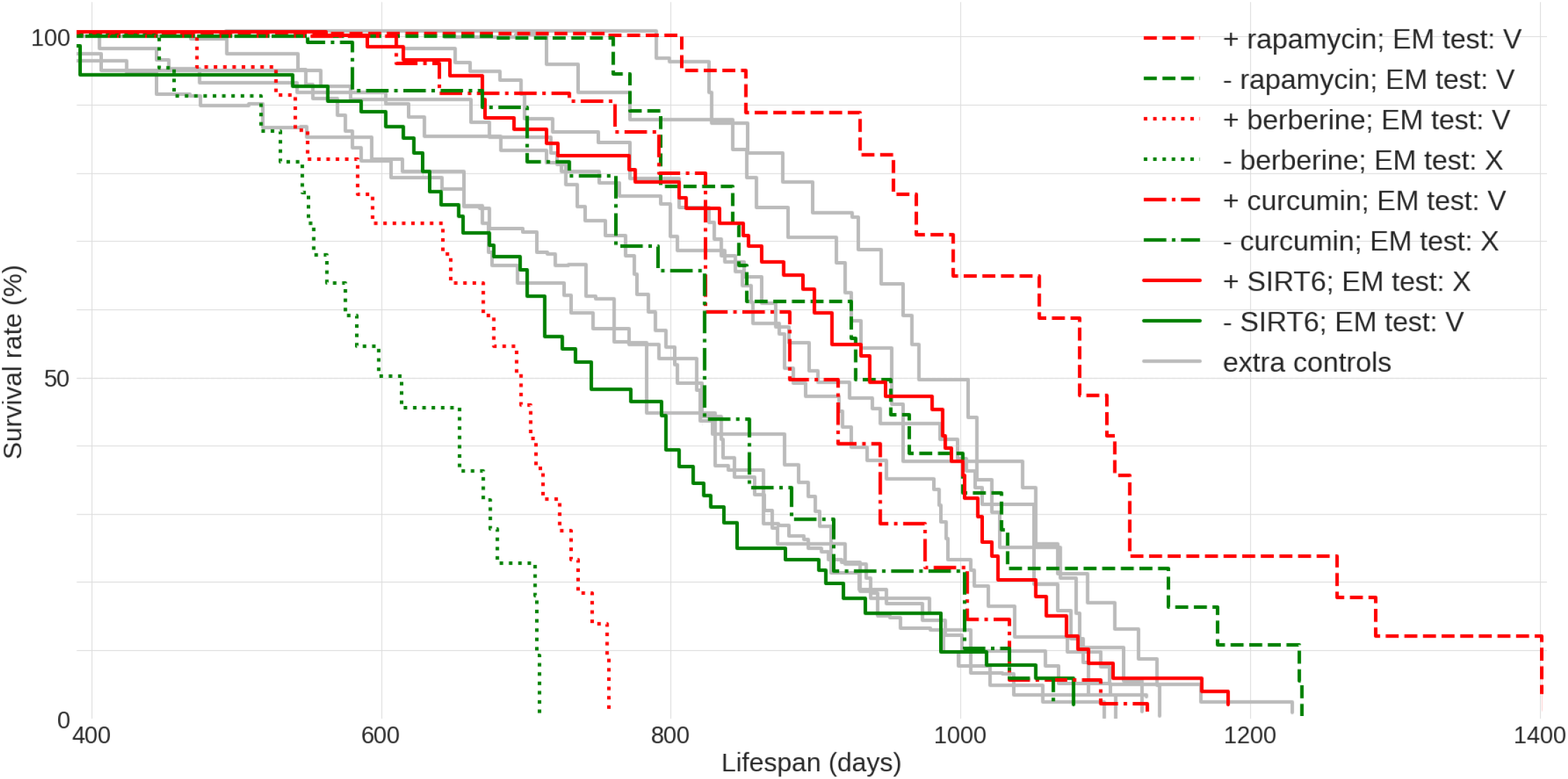
Lifespan data for C57BL/6J male mice from multiple lifespan extension and other unrelated studies. Four sample “successful” lifespan extension studies are shown for comparison: the control data is in green and respective intervention data is depicted via the matching stroke (solid or broken) in red. Each colored curve is labeled with its plausibility score P. The PMIDs of sources used here are: 27549339 (rapamycin), 31773901 berberine, 17516143 curcumin and 34050173 SIRT6. ‘Meta-control’ data given in gray here are replotted and annotated in Fig S1. EM - Extra Mortality test (see Supplement) was applied to datasets: V - passed, X - failed with the level of significance α = 0.01.

While compiling a comparison like that should be easy, we had to collect these data dealing with entirely different representations each time. It is hard to believe, yet there are no requirements nor data format standards today for submitting the lifespan data along with a publication. We believe that the scientific community needs to take action - to improve the quality of work and the reproducibility of scientific results within aging biology. To this end, we developed the lifespan data and meta-data standard (see Supplement). The format is simple yet flexible enough to allow comprehensive data description and re-analysis, we provide encoding for the berberine data from Fig.1 (see Supplement). Additionally, we have created a dedicated Web [resource](https://www.lifespan.io/animal-life-expectancy-comparisons/) “ALEC” (Animal Lifespan Expectancy Comparisons) for accumulation and interactive browsing of lifespan data. We plan to lobby the journals in the field and encourage the authors of previously published and forthcoming publications to adopt this format and to share the data using a central repository. Numerous large datasets of lifespan from studies in mice have already been loaded and are available for interactive browsing and comparison to a user-supplied lifespan data. This resource facilitates evaluation of new lifespan data via an instant validation in the context of present knowledge and testing the data with quality control techniques (see Supplement) specifically developed and implemented to ALEC. Unfortunately, our attempts to validate the lifespan improvements with other strains did not yield any qualitatively different results from the one we focus on in this study (data not shown).

## Discussion

In summary - in many studies reporting the lifespan extension of C57BL/6J mice, the lifespan of the intervention group appears significantly higher in comparison to the controls - yet is inferior to the lifespan of the control animals of the same strain reported elsewhere. To appreciate the significance of this point, we must consider that the primary practical motivation for experiments in biology of aging is to develop methods for extending the healthy lifespan. Naturally, such methods may differ from those needed for compensatory lifespan extension that is reduced because of genetic abnormalities, environmental challenges or suboptimal living conditions. Of note, the many interventions re-tested in C57BL/6J mice against deep phenotypes of health did not “slow aging” for most parameters monitored^11^. Thus, we posit that the majority of results in biology of aging may be irrelevant to the fundamental aim of this field and must be acknowledged appropriately.

## Supporting information

example of data entry

data format template

## Supplementary Materials and Methods

**Figure S1.**
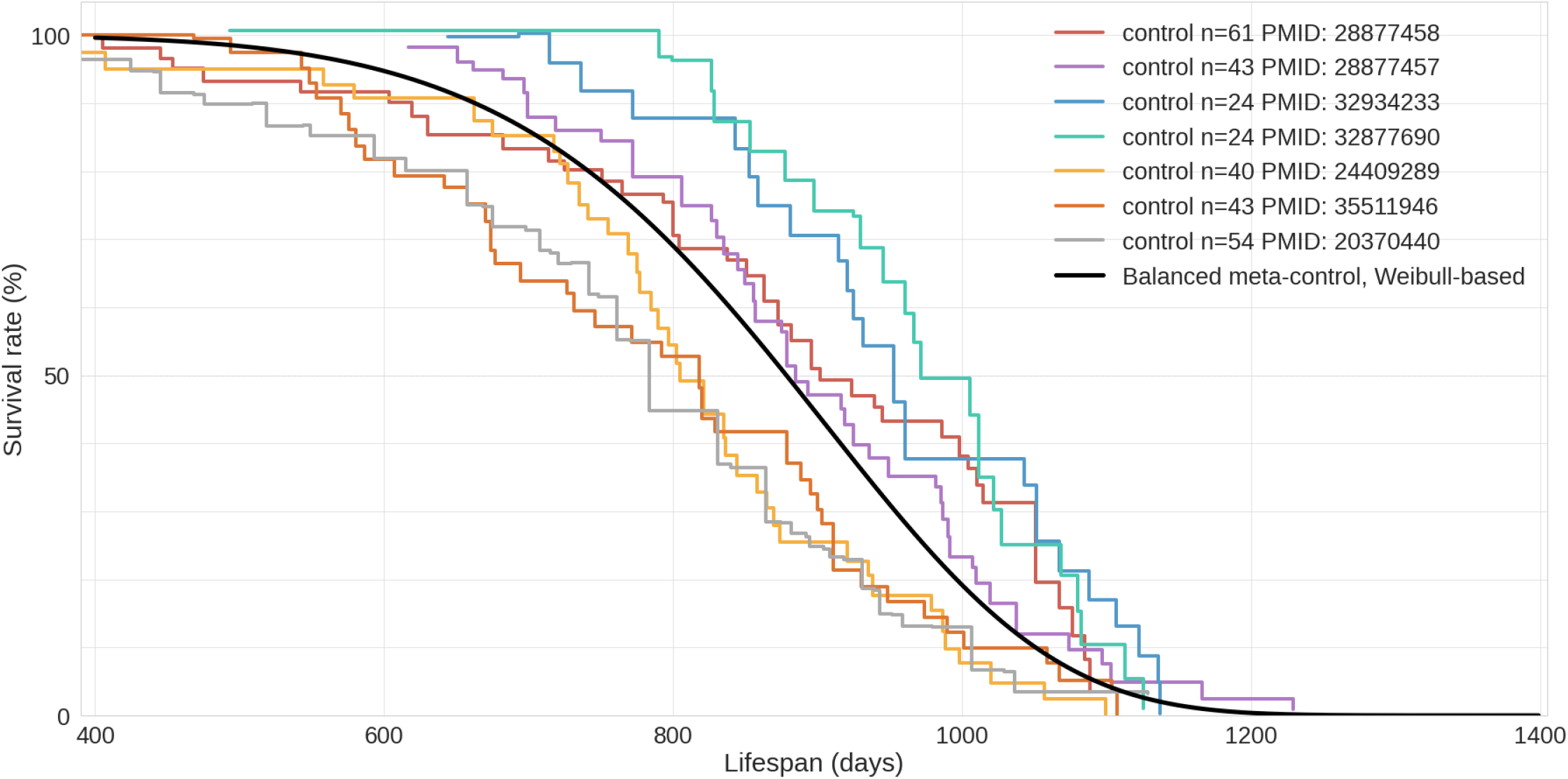
Selected meta-controls which were used as the background plotted in gray in the main text Figure 1 with respective source papers cited by PubMed IDs. Black solid line corresponds to a balanced meta-control constructed from parameters of fitted Weibull models of meta-controls (see Supplement).

### Lifespan Meta-data Format

The proposed data format is structured at three levels of hierarchy: a study meta-data, an experiment meta-data and a raw lifespan data. A given dataset is expected to only contain one study description, defining parameters specific to this study and common to all the experiments in a study (e.g. animal species used), where to find more information about the study (e.g. publication details and identity of whoever submitted the data). If a given dataset federates multiple studies, we expect the data to be broken up into multiple disjoint sets.

Each study is expected to contain several experiments, typically, at least a single intervention and a coupled control experiment but usually more, even though it is not impossible to imagine a single experiment aimed at collecting statistics ver normal lifespans. The experiment-specific meta-data defines what distinguishes a given experiment within this study from the others (e.g. specific drug used, gene knock out) as well as auxiliary data description.

Each raw data is trivially a set of numbers, one per line, each defining lifespan of a single individual. This minimalistic design allows for unambiguous data re-analysis. There are cases when additional per-individual data is desired, such as the health state of an animal, whether it was introduced separately or removed early, co-location, accidental injury. These can be provided in a separate CSV file ordered in the same way as the lifespan data, where specifics of the columns/fields are defined in the respective experiment meta-data “remarks” field.

We supply a set of Excel files to provide a template (three files: study, experiment, lifespan) for publishing new data and to illustrate (five files: study.xlsx, experiment1.xlsx, experiment2.xlsx, lifespan1.xlsx, lifespan2.xlsx) the format using one of the studies from Fig. 1.

### Intra-study extra-mortality test

To test the consistency of a given dataset we developed an extra-mortality test checking if the given dataset has an “unexpected” increase in mortality by comparison with two reasonable models of survival curves. To test this for a given dataset we first estimate a time derivative *dS/dt* of its survival function *S*. Next we estimate these derivatives from samples drawn from corresponding Weibull and Gompertz approximations of the dataset. To account for different measurement times (i.e. times when researchers recorded the number of died mice) for Weibull and Gompertz models, we used measurement times as in the original experiment, thus binning the observations similarly. By repeating sampling from model distributions 1000 times we compared maximal derivative *dS/dt* of original survival curve and model-sampled (Fig. S2, S3). If the maximal derivative in the original study is greater than one from 1000 model samples we consider such a dataset having the extra-mortality artifact. For example, “Sirtuin control” dataset doesn’t have extra-mortality by comparing with 1000 samples from the Weibull model of the dataset (Fig. S2, right event plot), and, thus, can be considered as of high quality. In contrast, “Berberine control” has such an artifact (Fig. S3, right event plot) and should be analyzed with caution.

### Inter-study plausibility test for a new control dataset

For testing the plausibility of a new control dataset we propose the following procedure. In the procedure we will rely on the comparison of a new dataset with selected meta-control datasets. We first assume that each meta-control dataset can be approximated with Weibull distribution of parameters λ and ρ which are scale and shape correspondingly. For fitting the Weibull distribution we use [WeibullFitter](https://lifelines.readthedocs.io/en/latest/fitters/univariate/WeibullFitter.html) class of [lifelines](https://lifelines.readthedocs.io/en/latest/index.html) package. WeibullFitter returns each of the parameters with corresponding standard error of estimate which can be used as intra-study variance for the parameter depending on a sample size and structure of a meta-control dataset. By fitting all meta-controls we obtain a set of parameters *λ* and *ρ* with corresponding standard errors of estimates.

We next assume that these parameters are drawn from a two-dimensional Gaussian distribution with unknown covariance matrix which we call reference distribution *P*(λ, ρ). Using the gathered estimates of parameters and their intra-study variances we estimate the covariance matrix and mean vector of the reference distribution. For that we calculate weighted mean, weighted marginal variances and weighted covariance of parameters using their inverse squared standard errors as weights and, thus, accounting differences in inter-study heterogeneity.

Once the reference distribution is computed we may use it for testing the “plausibility” of a new dataset. This can be achieved by estimating its parameters λ and ρ of corresponding Weibull fit and by computing the distance of the obtained vector of parameters to the corresponding mean vector of the reference distribution by simply applying Hotteling’s T-squared test. The result of the test is a value of statistics (squared Mahalanobis distance) and p-value. By accepting a reasonable value of significance α, this p-value can be used for testing a plausibility of a new dataset. If p-value is small - the dataset is implausible because of its dissimilarity with meta-controls.

### Balanced meta-control model for inter-study testing a treatment effect

Balanced meta-control is a hypothetical control model that harmonizes all accepted meta-controls in ALEC. For constructing balanced meta-control we first assume that each meta-control dataset can be approximated with Weibull distribution of parameters λ and ρ which are scale and shape correspondingly. For fitting the Weibull distribution we use [WeibullFitter](https://lifelines.readthedocs.io/en/latest/fitters/univariate/WeibullFitter.html) class of [lifelines](https://lifelines.readthedocs.io/en/latest/index.html) package. WeibullFitter returns each of the parameters with corresponding standard error of estimate which can be used as intra-study variance for the parameter depending on a sample size and structure of a meta-control dataset. By fitting all meta-controls we obtain a set of parameters λ and ρ with corresponding standard errors of estimates. For obtaining a balanced average of these parameters we use a meta-regression approach from [pymare](https://pymare.readthedocs.io/en/latest/) package (meta_regression function) for combining separate estimates with their standard errors. This approach returns balanced average parameters *λ* and *ρ* by accounting variances of Weibull estimates. The resulting parameters can be used for constructing meta-Weibull distribution which is the balanced meta-control. Next this balanced meta-control model can be used for computing the effect of a new treatment by, for example, comparison of median lifespans of a new survival dataset of treated mice with balanced meta-control median lifespan.

**Figure S2.**
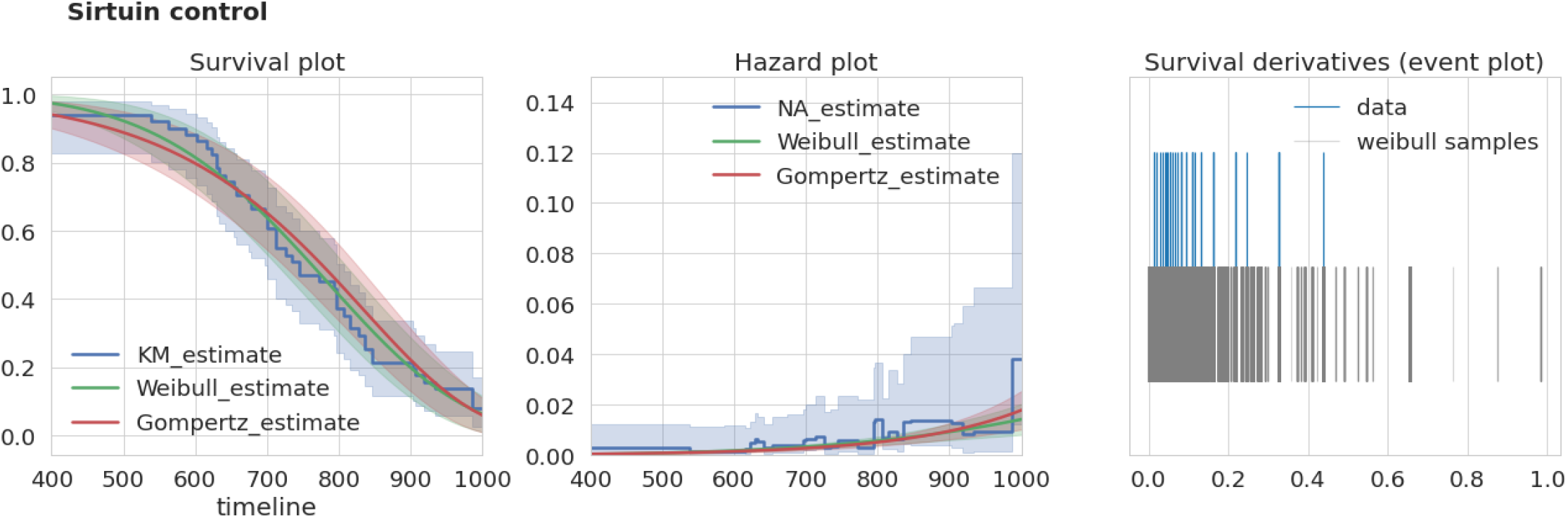
“Sirtuin control” dataset characteristics. Left, survival curves estimates from nonparametric and parametric estimators. Middle, corresponding hazard (mortality) estimates. Right, event plot (events distribution) of original data and 1000 samples of survival curve derivatives. KM - Kaplan-Meier estimator, NA - Nelson-Aalen estimator.

**Figure S3.**
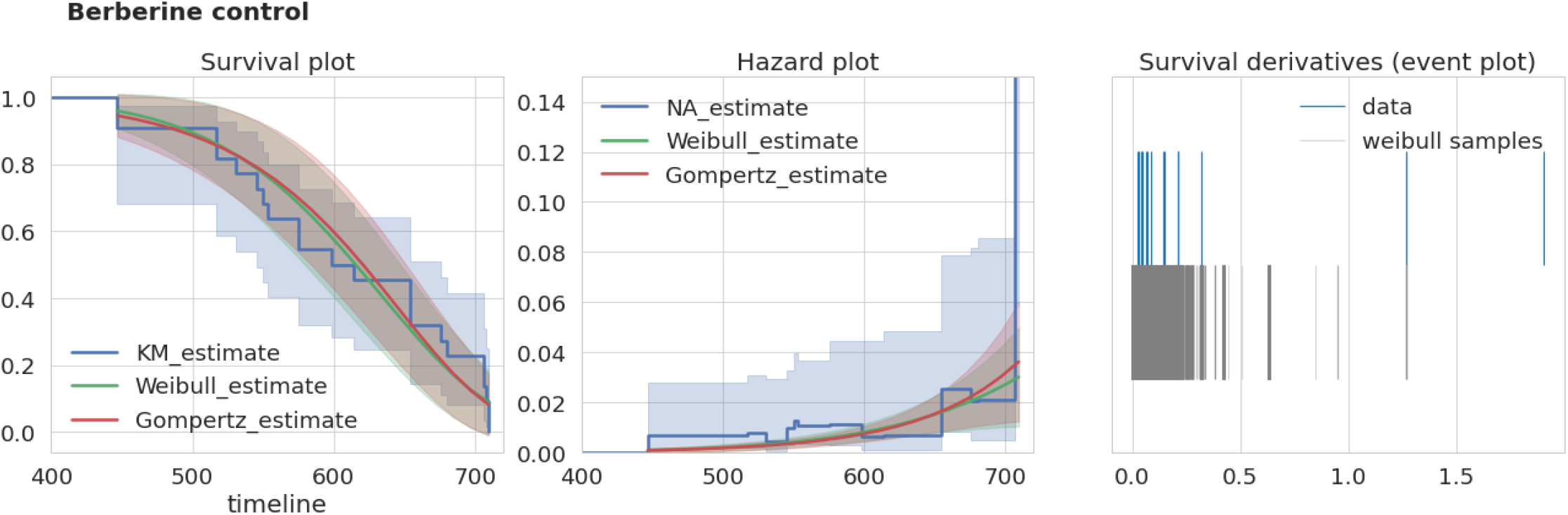
“Berberine control” dataset characteristics. Left, survival curves estimates from nonparametric and parametric estimators. Middle, corresponding hazard (mortality) estimates. Right, event plot (events distribution) of original data and 1000 samples of survival curve derivatives. KM - Kaplan-Meier estimator, NA - Nelson-Aalen estimator.

**Table S1.**
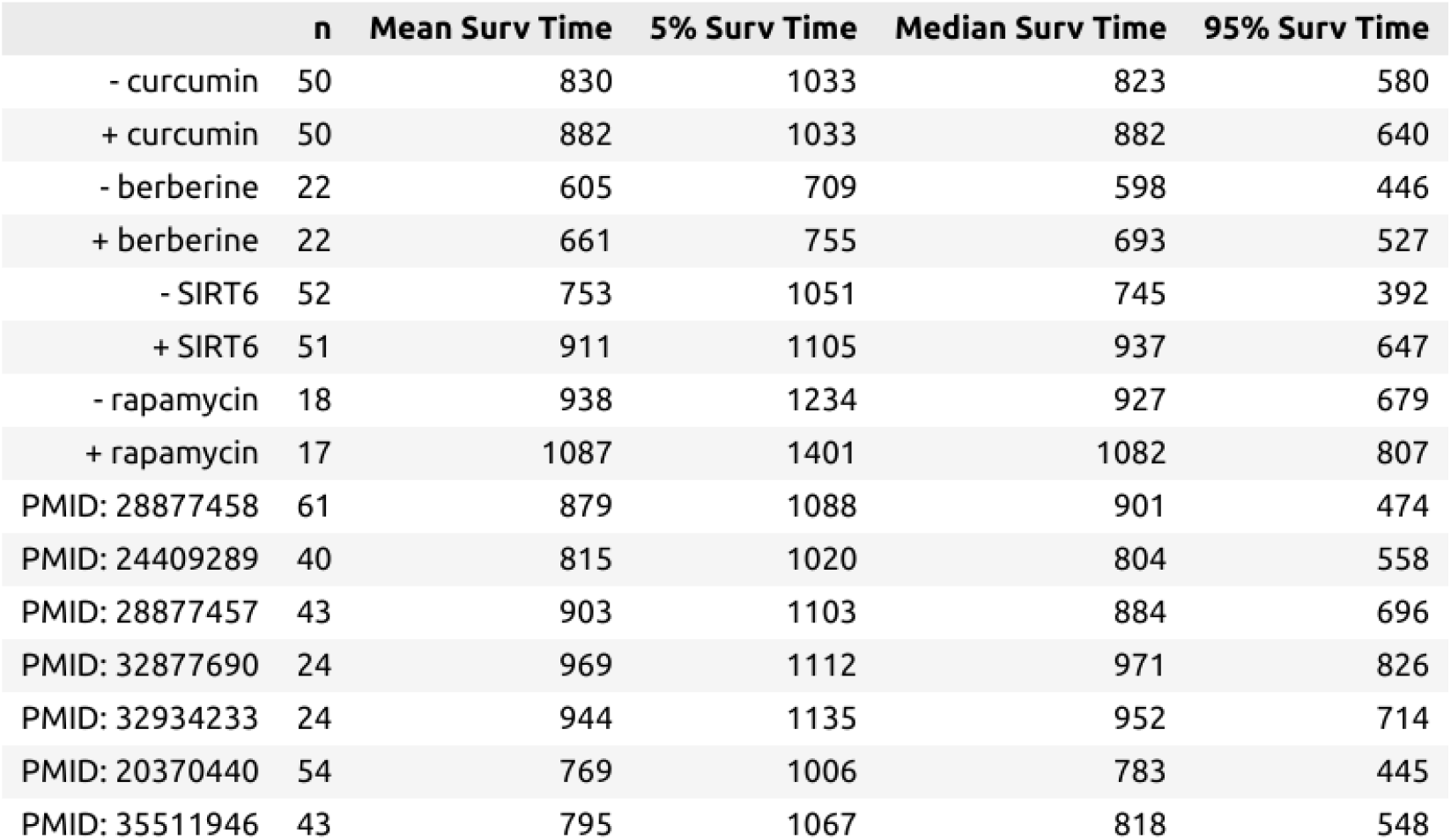
Characteristics of datasets from figure 1 of the main text.

